# Analysis of the optimality of the Standard Genetic Code

**DOI:** 10.1101/032748

**Authors:** Balaji Kumar, Supreet Saini

**Affiliations:** Department of Chemical Engineering, Indian Institute of Technology Bombay, Powai, Mumbai - 400 076, India

**Keywords:** Standard Genetic Code, Optimality, Frameshift, Point Mutation

## Abstract

Many theories have been proposed attempting to explain the origin of the genetic code. While strong reasons remain to believe that the genetic code evolved as a frozen accident, at least for the first few amino acids, other theories remain viable. In this work, we test the optimality of the standard genetic code against approximately 17 million genetic codes, and locate 18 which outperform the standard genetic code at the following three criteria: (a) robustness to point mutation; (b) robustness to frameshift mutation; and (c) ability to encode additional information in the coding region. We use a genetic algorithm to generate and score codes from different parts of the associated landscape, and are, as a result presumably more representative of the entire landscape. Our results show that while the genetic code is sub-optimal for robustness to frameshift mutation and the ability to encode additional information in the coding region, it is very strongly selected for robustness to point mutation. This coupled with the observation that the different performance indicator scores for a particular genetic code are seemingly negatively correlated, make the standard genetic code nearly optimal for the three criteria tested in this work.

## Introduction

Genetic code is an assignment of codons to amino acids, and defines the translational system for protein synthesis. Looking at the definition of a genetic code as a combinatorics problem, it equates to allotment of codons (say, non-identical balls) to amino acids and a stop signal (say, boxes). The total number of possible genetic codes therefore equates to solving the problem of number of ways to distribute the 64 non-identical balls among the 21 unique boxes, ensuring that each box gets at least one ball. A simple calculation shows that the total number of solutions available for this problem is of the order of 10^83^. The standard genetic code is only one of the possible solutions to this problem. What makes the standard genetic code so special against all other possible solutions? Or is the choice of standard genetic code being most used in all life forms random and a result of a chance event?

One of the key features of the standard genetic code is redundancy, where more than one codon corresponds to the same amino acid (1). As an example, in standard genetic code, Leucine is coded by six codons (UUA, UUG, CUU, CUC, CUA, and CUG). It is interesting to note that the nucleotide U, present in the 2^nd^ position is common for all the codons but only those in the first and third positions vary. Similarly, in the case of glutamic acid, there are two codons GAA and GAG, for which only the nucleotide located at the 3^rd^ position differs. The effect of redundancy is that, degeneracy in the third position of the triplet codon cause only a silent mutation i.e. there is no effect of mutation in protein translation because the biochemical property is conserved by equivalent substitution of amino acids.

One of the theories for the evolution of the standard genetic code is that it is thought to be a frozen accident during evolution. This theory states that “genetic code is a random, highly improbable combination of its components formed by an abiotic route, and altering it from its present state would be disadvantageous”, which implies that the mechanism of allocating codons to amino acids is entirely a matter of chance (2). Several recent studies however suggest that the genetic code is not a frozen accident but has evolved so as to minimize transcriptional and translational errors (3). It has been shown that the standard genetic code minimizes the effect of point mutations or mistranslations: either the erroneous codon is a synonym of the original amino acid, or it encodes an amino acid with similar biochemical properties (4).

Standard genetic code, when compared with a truly random code, was observed to be partially optimized for robustness to frame-shift and point mutations (5). Canonical genetic code outperforms generated random codes in terms of polar requirement scale (6, 7), where polar requirement is the biochemical property of each amino acid defined by the paper chromatography experiments of Woese and co-workers (8).

The standard genetic code has also been shown to be capable of including additional information within protein-coding sequences (9). These additional data can be biological signals like binding sequences for regulatory and structural proteins (10-12), and splicing signals that include specific 6–8 base pair sequences within coding regions and mRNA secondary structure signals (13-15). For comparison of standard genetic code with other possible codes, simulations have been performed in the past. However, the computationally intractable large number of possible genetic codes means that only a miniscule fraction of all possible genetic codes can be analyzed. For example, in a study by Koonin and co-workers (5) rules were defined to limit the possible number of genetic codes, and a small fraction of the possible codes were then generated and analyzed.

Some of the other attempts in this regard have also been carried out recently (16, 17). Schonauer and Clote iterate over millions of codes to explore the optimality of the genetic code in a much larger space and mapping of {‘A’, ‘T’, ‘C’, ‘G’} onto the 20 amino acids and one stop codon. Sergey Naumenko et al and Churchill et al., 1990 discuss the importance of stop codons on the optimality of the standard genetic code, and suggest that among all genetic code mark-ups with three stop codons, the standard genetic code mark-up has the maximum possible probability of the terminating the mis-translation process (16-18).

In this work, we try and analyse the optimality of the standard genetic code against randomly generated genetic codes across three performance criteria: (a) robustness to point mutations, (b) robustness to frameshift mutations, and (c) the ability of a code to encode additional information in the coding region. We first present results from a local sampling from the “sequence space” associated with the genetic codes, and compute scores across the three indices. We show that in this local sampling, the standard genetic code comes out as an almost optimal genetic code. However, our results show that the same trends/qualitative features of the standard genetic code do not all hold up when the genetic code sampling is more diverse from its “sequence space”. Our results indicate that the genetic code is sub-optimal for robustness to frameshift mutation and ability to encode additional information; it is strongly selected for robustness to point mutations. Last, we note that the performance of a genetic code across the difference indicators seems to be negatively correlated.

## Methods

To analyse the optimality of the standard genetic code, we use the following performance parameters: robustness to frame-shift and point mutations; and the ability to encode additional information in the coding region of the genome of *Escherichia coli* (str. K 12 substr. DH10B chromosome) *(E. coli)*. The length of the genome is 4.6 million bases (18). Computations and analyses were done using Perl and Python.

### Generation of Random Genetic Codes

To compare the performance of the standard genetic code against other possible genetic codes, random genetic codes were generated using Perl. The randomly generated codes were designed based on three criterion as follows. Within each criteria, 10,000 codes were generated randomly.

**Random Codes 1 (RC1)**. 10,000 genetic codes were generated by random allocation of codons to amino acids while ensuring that the number of codons allotted to each amino acid (and stop signal) is the same as that in the standard genetic code.

**Random Codes 2 (RC2)** In this case, 10,000 genetic codes which satisfy the following two properties were generated. First, as in RC1, the number of codons allotted to each amino acid was same as that in standard genetic code. Second, codon allocation was done in a semi-random manner, where only codons which correspond to polar amino acids in the standard genetic code were re-allocated between polar amino acids (and codons corresponding to non-polar amino acids were re-allotted to non-polar amino acids only). This was done to ensure that the localized structure of biochemical properties in the genetic code is preserved. In this set, the codons corresponding to the stop codons were kept the same as that in standard genetic code.

**Random Codes 3 (RC3)** An identifying feature of the standard genetic code is its “block structure” where all codons allocated to an amino acid occur as a “block”. To preserve this structure, 10,000 genetic codes were generated ensuring that this structure of the standard genetic code is preserved. For allocation of stop codons, it was ensured that two of the three stop codons differ only in the third position, and that the third stop codon differs in the second position (just as in the standard genetic code).

### Genetic Algorithm

To generate genetic codes with performance better than that of standard genetic code, we implemented a genetic algorithm with three separate fitness functions. The fitness functions namely optimized the point mutational robustness, frameshift robustness, and ability to encode parallel information, in the genetic code. In this algorithm, we started with a population of nineteen randomly generated codes, and the standard genetic code to kick-start the evolution. The population size was maintained constant at 20. In each generation of the simulation, genetic codes were mutated, recombined, scored for their fitness and the fitter codes were selected for the next generation. The probability of a mutation was defined as the chance that a codon assigned to an amino acid is re-assigned to another amino acid chosen randomly. This value was taken to be 0.05. The mutation rate was set at 0.1, meaning approximately two codes undergo mutation every generation on average. The 64 codons in the genetic code were numbered from 1 to 64. Recombination between the two codes was defined at codon number X, such that all codons with numbers less than X are taken from code I, and all codons from numbers X to 64 are taken from code II. Two codes were chosen and recombined randomly in each generation. After mutation and recombination, the viability (that all 20 amino acids and stop signal were represented in the code) of the new “evolved” codes was verified. The new genetic codes were then scored based on fitness scoring as follows, and the fitter ones were selected to the next generation, based on roulette wheel sampling.

### Quantification of performance of genetic codes

#### Point mutational Robustness

Amino acids were grouped based on their biochemical property into:

- <u>Non-Polar</u>: glycine (Gly), alanine (Ala), valine (Val), leucine (Leu), isoleucine (Ile), proline (Pro), phenylalanine (Phe), methionine (Met), and tryptophan (Trp).
- <u>Polar-uncharged</u>: serine (Ser), threonine (Thr), cysteine (Cys), asparagine (Asn), glutamine (Gln), and tyrosine (Tyr).
- <u>Acidic</u>: aspartate (Asp) and glutamate (Glu).
- <u>Basic</u>: arginine (Arg), lysine (Lys), and histidine (His)

Point mutational scoring system takes into account (a) biochemical property of amino acids, and (b) relative sizes of amino acids. Every point mutation belongs to one of the following three: (a) silent - no change in amino acid, (b) conservative - amino acid mutates to a biochemically similar amino acid, and (c) non-conservative. A scoring system for each code was implemented for each of the 576 mutations – each codon mutated 9 possible times. If a mutation belonged to (a) one point was awarded, if it belonged to (b) 0.5 was awarded, and no points were awarded for (c). Additionally, amino acids were ranked from smallest to largest amino acid by size (using molecular weight as proxy). The fraction of size conserved, or fraction of size changed subtracted from unity, was also added to the score of a codon. This was done only for cases excluding the stop codon. The biochemical property and amino acid sizes were given equal weights. The cumulative score is the score of a genetic code. All generated genetic codes were scored similarly.

#### Frameshift robustness

Second, to search for codes better at frame-shift robustness an altered genetic algorithm was devised. A fitness function was implemented which quantifies the probability with which a faulty peptide translation will be terminated, taking into account the amino acid frequencies of *E.coli*.

We calculate the theoretical probability of encountering a stop in a misread frame, by using di-codon sequences (9). We consider all 61x61 combinations of codons, excluding the three stop codons. Stop is encountered in 2-4 position for +1 frame shift and 3-5 position for - 1frame shift. Probability of encountering a Stop codon in an insertion frame is the sum of all probabilities of di-codons with Stop in 3-5 positions. Similarly, probability of encountering a Stop codon in a deletion frame is the sum of all probabilities of di-codons with Stop in 2-4 positions. Probability of a di-codon sequence is calculated as follows. A codon C coding for an amino acid A, occurs with a probability of frequency(A)/(Number of synonymous codons of A). Probability of a di-codon is product of probabilities of the two codons. Here, we assume uniform codon-usage for ease of calculations, without compromising on the accuracy of the scoring systems.

#### Parallel coding ability

Here we calculate the probability of encoding N-base sequences in the coding regions of E. *coli* (9). We considered a value of five for N in this work. We take the fitness score of a code as the probability to encode its top 20% most difficult N-base sequences or N-mers (for N=5). Probability of each 5-mer is the combined probability with which it can be incorporated in three reading frames - correct Open Reading Frame, insertion, and deletion reading frames. In each frame, probability of a 5-mer is the sum of probabilities of all possible codons with which it can occur (See above for probability of codon occurrence).

## Results

### The standard genetic code is nearly optimal at minimizing point mutational errors

To start the analysis, we generated 30,000 genetic codes (10,000 each belonging to the group RC1, RC2, and RC3), and analyzed their performance by a point mutational scoring system (see methods section for more details on details of RC1, RC2, and RC3 codes; and the scoring system used). From our analysis, we note that upon introduction of a point mutation, the standard genetic code leads to minimum number of cases, where an amino acid is maximally replaced with another one. As shown in **Table 1**, a majority of the times, an amino acid is replaced by itself, after a point mutation. In addition, even if a point mutation was to lead to a change in the amino acid, the standard genetic code leads to maximal replacements such that the biochemical properties of the amino acid are conserved. Among the 30,000 codes we tested in this section only 38 genetic codes outperformed the standard code with respect to their resistance to change in amino acids as a result of point mutations. This indicates that the standard genetic code is nearly optimal for minimizing the point mutational errors.

**Table 1:** Analysis of point mutations to the codons encoding for all twenty amino acids (column 1). The columns 2-5 give the amino acid encoded most often after introduction of point mutation to each amino acid.

| Amino acids | Standard | RC1 | RC2 | RC3 |
| --- | --- | --- | --- | --- |
| Alanine | Alanine | Leucine | Leucine | Proline |
| Arginine | Arginine | Leucine | Leucine | Leucine |
| Asparagine | Lysine | Serine | Threonine | Serine |
| Aspartate | Glutamate | Arginine | Serine | Arginine |
| Cysteine | Arginine | Arginine | Leucine | Arginine |
| Glutamate | Aspartate | Arginine | Arginine | Arginine |
| Glutamine | Serine | Arginine | Serine | Threonine |
| Glycine | Glycine | Leucine | Leucine | Leucine |
| Histidine | Glutamine | Arginine | Leucine | Arginine |
| Isoleucine | Isoleucine | Arginine | Lysine | Leucine |
| Leucine | Leucine | Arginine | Arginine | Isoleucine |
| Lysine | Asparagine | Leucine | Isoleucine | Arginine |
| Methionine | Isoleucine | Isoleucine | Leucine | Leucine |
| Phenylalanine | Leucine | Leucine | Arginine | Aspartate |
| Proline | Proline | Arginine | Leucine | Alanine |
| Serine | Serine | Arginine | Arginine | Threonine |
| Threonine | Threonine | Arginine | Leucine | Serine |
| Tryptophan | Arginine | Isoleucine | Arginine | Glycine |
| Tyrosine | Tyrosine | Arginine | Arginine | Serine |
| Valine | Valine | Arginine | Leucine | Leucine |

### Standard Genetic Code is nearly optimal at minimizing frameshift errors

Next we compared the performance of the 30,000 genetic codes with that of the standard genetic code at minimizing frameshift errors. The genetic codes were scored by introduction of a frameshift mutation, and noting the number of amino acids that are added to the faulty peptide chain before the ribosome encounters a stop codon. The score is inversely proportional to the length of this peptide chain. In our analysis, we note that of the 30,000 codes tested only 84 outperformed the standard genetic code (2 in RC1, 2 in RC3, and 80 in RC3). This corresponds to the standard genetic code outperforming 99.72% of all codes in the three groups at frameshift error minimization.

In a previous work (9), the ability of the standard genetic code to be nearly optimal at frameshift robustness was attributed to the allocation of stop codons. Upon generating all genetic codes with three stop codons (but with the wobble constraint), we note that the standard genetic code is nearly optimal among 5472 codes (including the standard genetic code) generated this way. In this analysis, 61 of all the codes outperformed the standard genetic code at frameshift robustness.

In the same work, Alon and coworkers show that the standard genetic code is also optimal for encoding additional information in the coding regions of the genomes. This additional information is thought to include: (a) binding sites for regulatory proteins that bind coding region (10-12, 19); (b) DNA and mRNA binding proteins (20); (c) histones binding sites (21-23); (d) Splicing signals (24); and (e) mRNA secondary structure signals (13-15, 25). Testing the ability of the standard genetic code to encode additional information in the coding sequence and its robustness to frameshift mutation against all 5472 codes, we note that the standard genetic code is nearly optimal for these two features <u>(**Figure 1**)</u>. Here, the addition information encoding ability is quantified as the average probability of encoding an N-base sequence (N=6 and averaged over all 4^^^6 = 4096 sequences). The proteome considered, was average amino acid frequencies from 134 organisms as previously reported.

**Figure 1.**
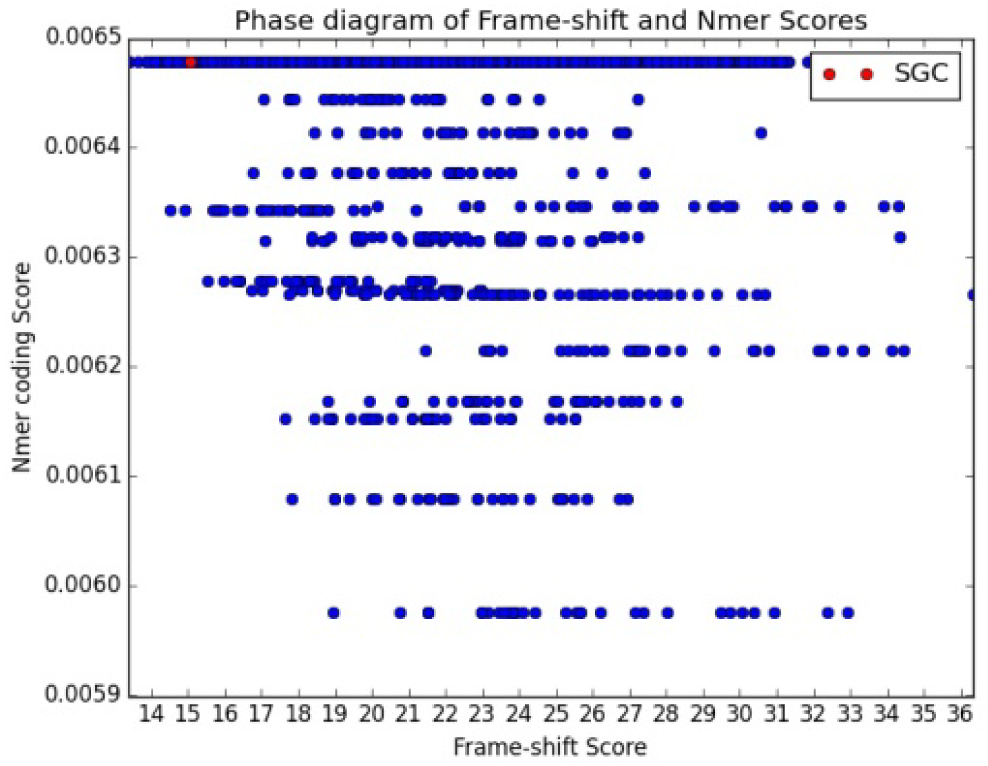
Performance of 5472 codes (from shuffling Stop codons with Wobble constraint) in frameshift robustness and additional information encoding ability. Expected faulty peptide length before termination and average probability of encoding an N-mer of size 6 are plotted. Amino acid profiles taken into consideration were averaged from 134 organisms.

However, we note that the standard genetic code is average at encoding additional information in the coding sequences, when the ease of a genetic code to encode the most difficult X-percent of the N-mers in the coding region is analyzed. We note that for both N = 5 and N = 6, the standard genetic code performs around the average for the most difficult 5% N-mers, among the 5472 codes (**Figure 2**). These results hold independent of the choice of “most difficult X%”, as shown in **Figure Supplement 1**.

**Figure 2.**
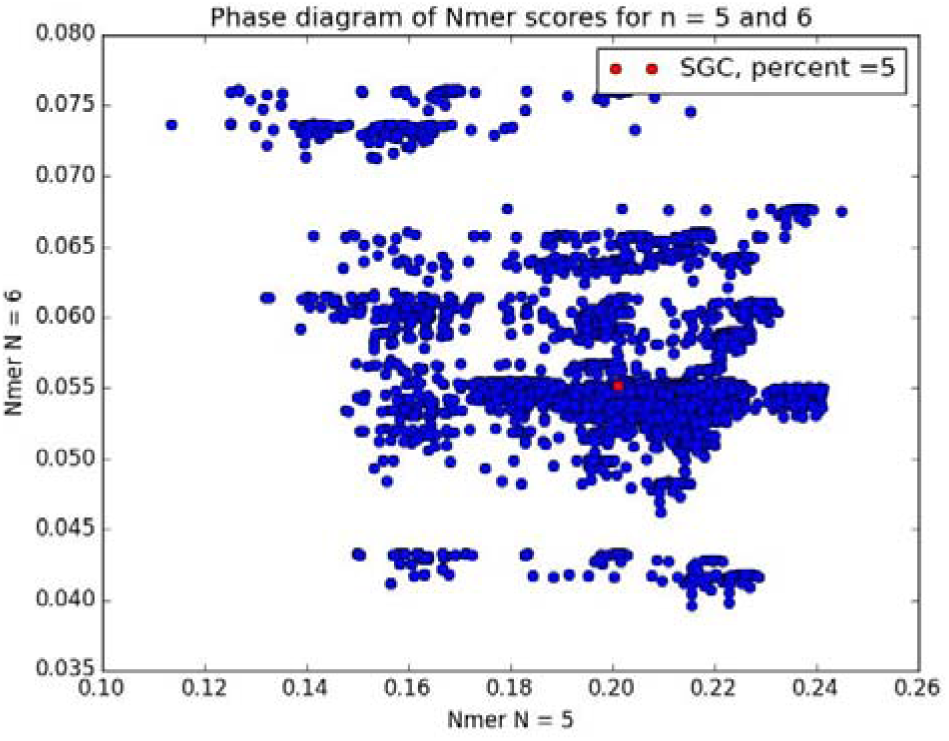
Performance of the 5472 codes at encoding additional information, when different parameters are considered. For each code, probability of encoding its top X% most difficult N-mers is plotted for N = 5 and N = 5, and X = 5

As a result of these conflicting conclusions regarding the optimality of the genetic code, we developed a genetic algorithm to scan a much larger pool of genetic codes, and compare the performance. This was done to ensure that the genetic codes being analyzed were from different sections of the fitness landscape associated with the sequence space corresponding 10^83^ codes, and that the fitness landscapes, in general, tend to be rugged with multiple peaks (26). By scanning a very small sub-set of these genetic codes in a systematic manner, we were likely only scanning a small, biased set of genetic codes. This set, we speculate, is not representative of the entire space defined by all genetic codes, and hence, our choice to use a genetic algorithm.

### Search for genetic codes which out-perform the Standard Genetic Code

#### Robustness to point mutational load

We first used the genetic algorithm to search for codes that can minimize point mutational errors better than standard genetic code. We implemented a scoring system that takes into account the biochemical property and the size of an amino acid, as these two properties play key roles in dictating protein functionality (see methods for more details). Through the genetic algorithm, we sampled approximately 15 million distinct genetic codes. Among these, we were specifically interested in those genetic codes with scores more than 616.26, which corresponds to the score of the standard genetic code at point mutational load minimization. Among all the codes scanned, only 64 genetic codes were found that outperformed the standard genetic code at this feature. This set of genetic codes had scores ranging from 616.39 to 635, and hence, outscored the standard genetic code by less than five percent. The distribution of scores for codes which outperformed the standard genetic code is as shown in **Figure 3**. As shown, among these codes, a majority are better than the standard genetic code by less than one percent. The performance of standard genetic code was found to be statistically significant and highly optimal when compared to other possible theoretical codes (P = 1.22e-5).

**Figure 3.**
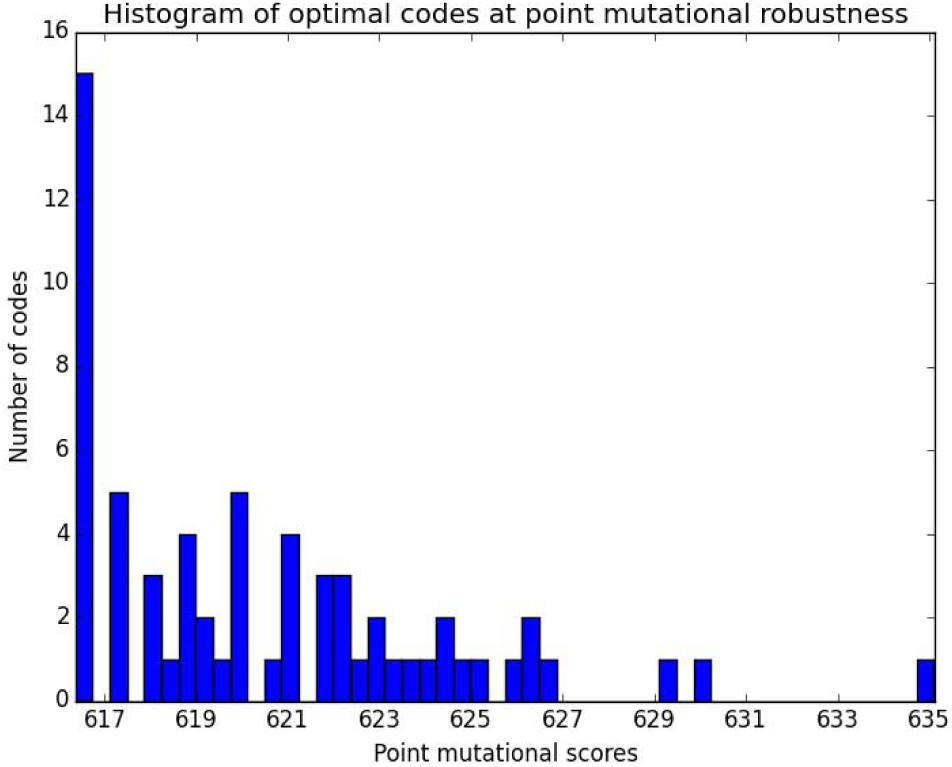
Histogram Plot of point mutational scores of genetic codes found, which outperformed Standard Genetic Code at the Genetic Algorithm with point mutational scoring fitness function.

#### Standard genetic code is sub-optimal but non-random at minimizing frameshift errors

Next, we used the genetic algorithm to score genetic codes for their ability to minimize frameshift errors (See methods). Our analysis with codes RC1, RC2, and RC3 indicates that the standard genetic code is selected for minimizing frameshift mutational errors. Similar results were found when we generated codes by randomizing the stop codons where only 1.1% percent of the codes out-performed the standard genetic code.

To search for codes with better scores at frameshift robustness than the standard genetic code, we used a genetic algorithm (with a modified fitness function as compared to the last section, see methods). Upon scanning approximately 1.6 million codes, we note that more than 90% codes (about 1.5 million) out-perform the standard genetic code. While the standard genetic code scores 0.062 in our scheme, of all the genetic codes analyzed, the mean score was about 0.22, and the highest 0.34. The distribution associated with the codes is as shown in **Figure 4**. Contrary to previous reports (9), and our analysis with the RC1, RC2, and RC3 codes, these results show that the standard genetic code is sub-optimal for robustness to frameshift mutations. Our results indicate that robustness to frameshift mutation has not been specifically selected for. We speculate that the possible reason(s) for this could be because frameshift errors are likely to only have a small effect on cellular fitness, as faulty peptides will simply be broken down by proteases before causing harm; in addition, translational errors are an order of magnitudes higher than transcriptional errors (27), because they allow faster sampling and hence evolution of proteins, without compromising the DNA. However, regardless, these results indicate the significance of scanning different regions of the fitness landscape associated with the genetic code space. While a local scan of this region might show the local optimality of the standard genetic code, a more comprehensive search of the landscape throws up totally different features/results.

**Figure 4.**
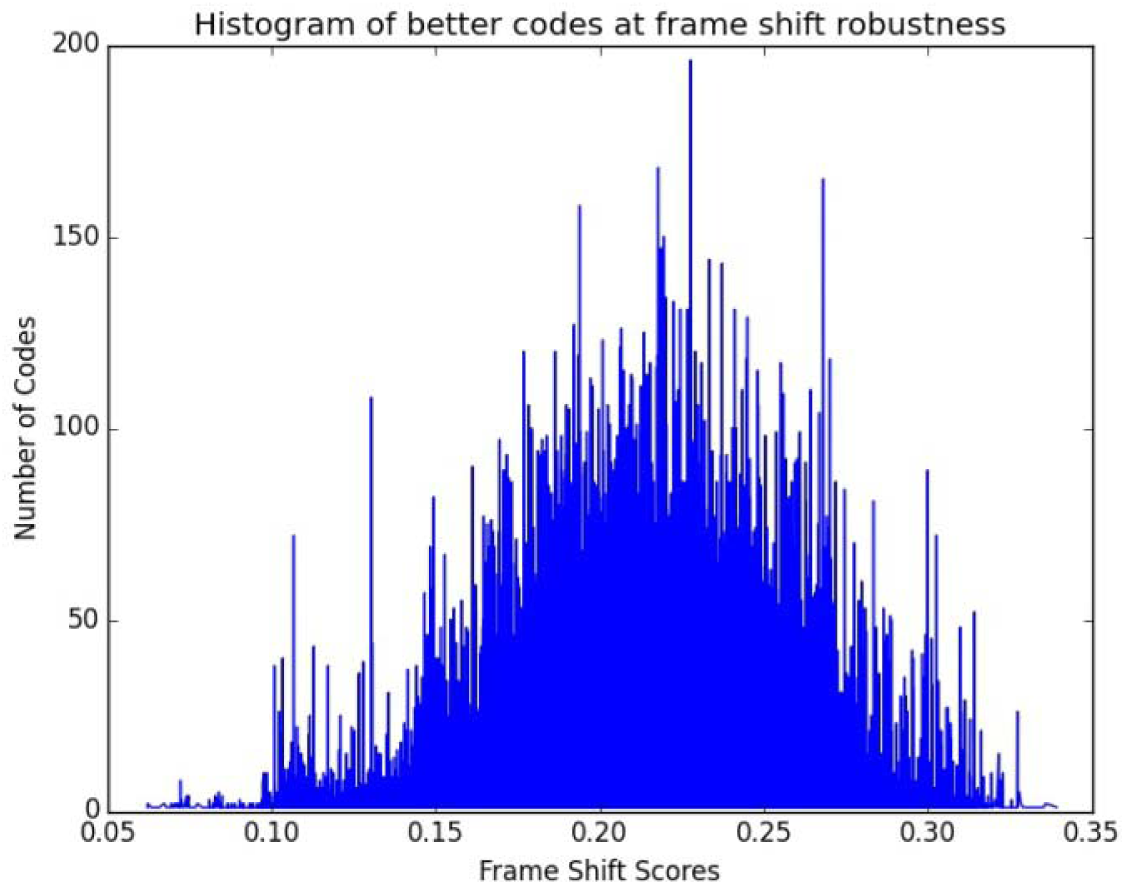
Histogram Plot of frameshift robustness scores of genetic codes found which outperformed Standard Genetic Code at the Genetic Algorithm with frameshift robustness fitness function.

#### Standard Genetic Code is sub-optimal at encoding additional information

Our analysis with shuffling of the stop codons shows that the standard genetic code is average at incorporating additional information in the coding region. This was independent of the percent of the most difficult N-mers that was taken for scoring, and also independent of the length of the N-mer (for both N = 5, and N = 6). We next used the genetic algorithm to scan parts of the sequence space which outperforms the standard genetic code at encoding additional information. For this purpose, we used the scoring system for N-mers of length five.

The standard genetic code was able to outperform roughly six percent of all genetic codes tested in our algorithm. Of the roughly 105,000 genetic codes tested, the standard genetic code fared worse off than 99,000 of these codes. The distribution of score among these codes represents a normal curve, and the standard genetic code lies at one end of this distribution (**Figure 5**). Our results show that there are many (in fact, most) locations in the sequence space where the genetic codes outperform the standard genetic code at encoding additional information in the coding regions of the genome. While previous results show that the standard genetic code might be a local optimum, but globally many peaks exist, and most of them perform better than the standard genetic code.

**Figure 5.**
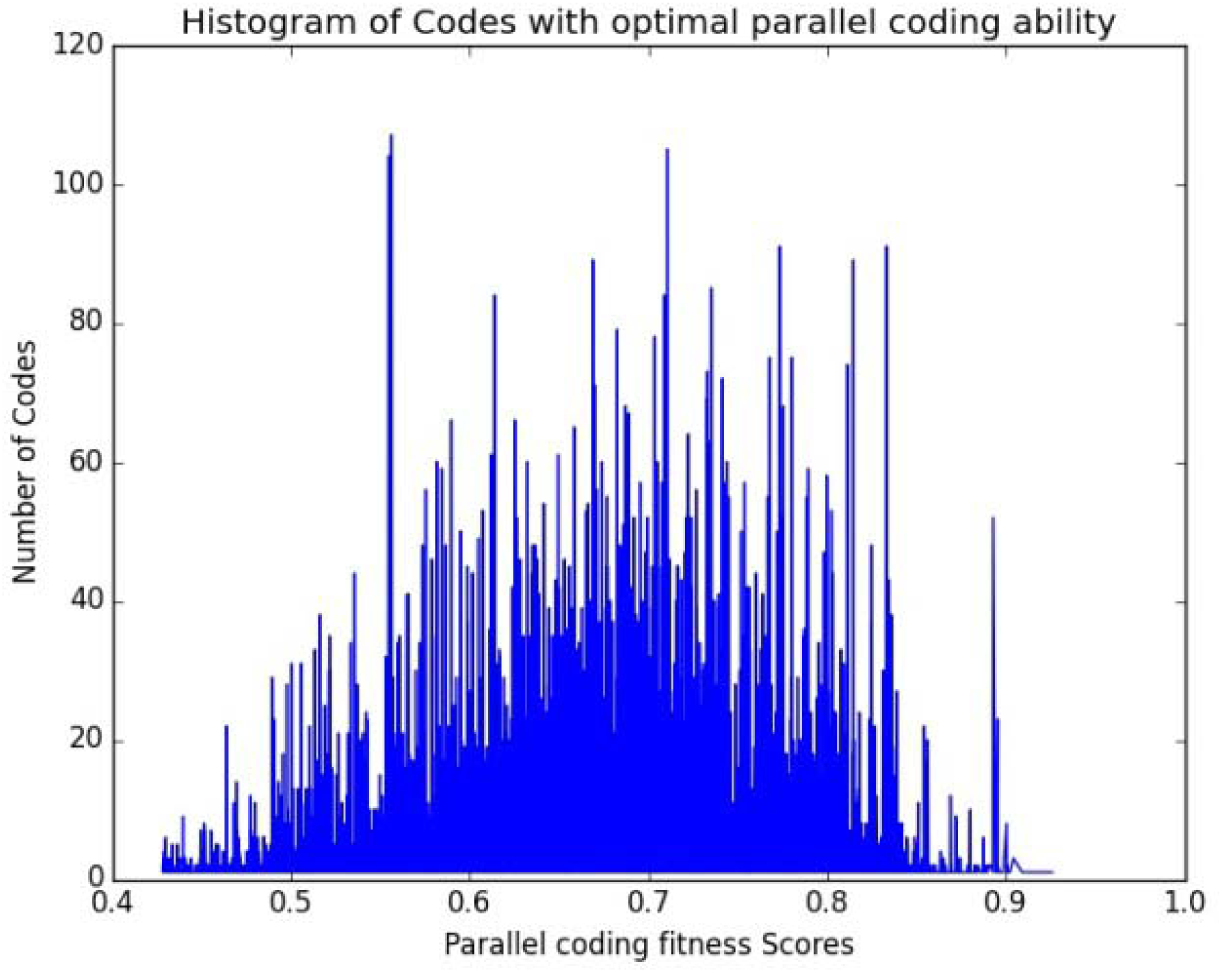
Histogram Plot of parallel coding ability of genetic codes found which outperformed Standard Genetic Code at the Genetic Algorithm with parallel coding ability fitness function.

### Robustness to frameshift mutation, robustness to point mutation, and ability to encode additional information – taken together, the standard genetic code is significantly better than other genetic codes

Lastly, we compared the performance of the genetic codes that outperformed the standard genetic code on any one of the three indices, by testing on the other two indices. For instance, we scored all 64 codes that outperformed the standard genetic code on their robustness to point mutations on their robustness to frameshift mutation and their ability to encode N-mers. The same process was followed for the other two scoring indices. Four codes out of 64 were found to outperform the standard genetic code on the other two indices as well. On the other hand, no genetic code from the other two sets outperformed the standard genetic code on all three scoring systems. Thus, of a total of roughly 16.9 million genetic codes tested, 4 outperformed the standard genetic code on all three indices tested here. The fact that only four out of 64 (only about six percent) were able to outperform the standard genetic code (on robustness to frameshift mutation and ability to encode additional information) is likely significant since in a random sampling of codes by the genetic algorithm, 90% outperform the standard genetic code. However, when selecting for genetic codes which outperform the standard genetic code at robustness for point mutation, and checking for the score of the selected codes for their ability to robustness to frameshift mutation and encode additional information, only about 6% of the codes satisfy the criteria. Thus, these results appear to indicate a trade-off while optimizing performance for multiple criteria.

This trade-off was again observed in a preliminary analysis when comparing performance of codes for their ability to encode information and robustness to frameshift mutation. On sampling a small set of 5,000 randomly generated codes (**Figure 6**), we note two distinct features: (a) there is a statistically significant inverse correlation between the ability of the code to encode additional information and its robustness to frameshift mutation, and (b) randomly generated codes form clusters in a plain indicating performance across the two criteria. The significance and the evolutionary relevance of this trade-off between performance criteria for genetic codes are being currently explored.

**Figure 6.**
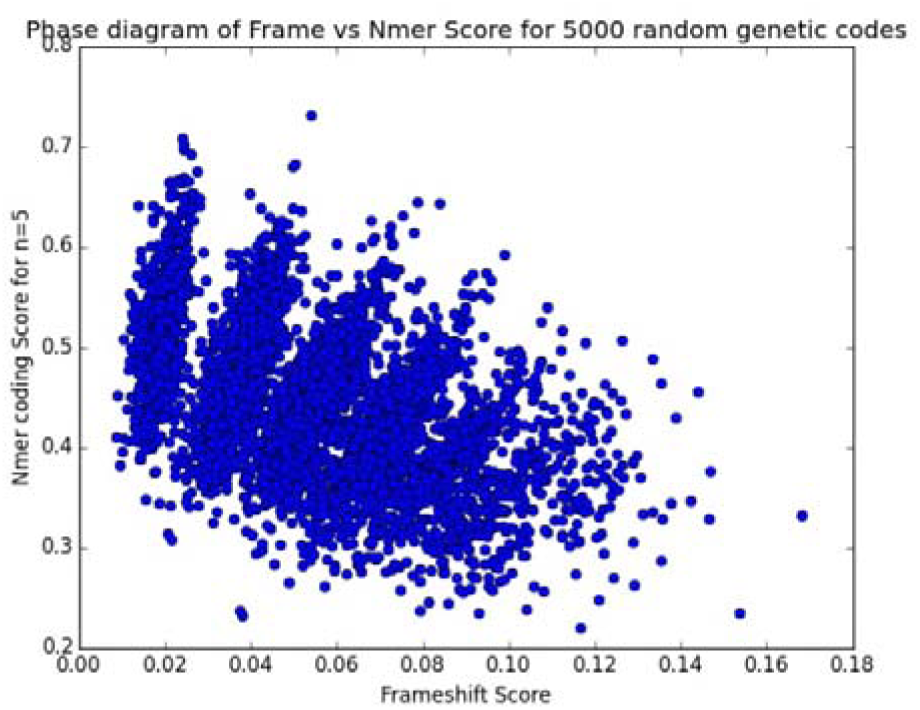
Performance of 5000 randomly generated genetic codes at encoding additional information (N = 5) and robustness to frameshift mutations.

On running the genetic code with all three objective functions combined into one, our preliminary scan of 52,000 genetic codes resulted in identification of 14 codes which outperform the standard genetic codes at all three performance indicators. Interestingly, multiple runs of the genetic algorithm lead to 186 genetic codes which outperform the standard genetic code. However, this group contained only 14 unique genetic codes (the rest being repeats). We are currently exploring (a) as to why the genetic algorithm leads to so many repeated solutions when optimized for all three performance indices, and (b) what is the allocation of codons to amino acid pattern in genetic codes which outperform the standard genetic code.

## Conclusions

In this work, we analyze the optimality of the standard genetic code across three features: (1) ability to truncate translation in case of frameshift, (2) ability to resist change in amino acid in case of a point mutation, and (3) ability to encode additional information in coding sequences. Our simulations suggest that the genetic code is nearly optimal in performance across these three criteria. However, looking at individual performance indicators, our results demonstrate that the standard genetic code is sub-average when compared randomly across codes for robustness to frameshift mutation, and ability to encode additional information. In fact, the near-optimality of the standard genetic code was observed largely due to its performance at robustness to point mutations. We also present some preliminary evidence for trade-off for a genetic code between the different performance criteria.

The performance of the genetic code likely was optimized across a number of criteria – like ability to incorporate or avoid short sequences in the coding region, control mRNA stability and structure, translational rates (9). Work in this direction has shown that the standard code performs close to the optimal level, when compared with other randomly generated codes. Presumably, the translation machinery emerged first for the amino acids which were synthesized first in the primitive atmosphere (28). How the later amino acids integrated into this translation machinery, giving shape to the fitness landscape associated with codon-amino acid assignment space, to produce an “optimal” genetic code remains an open question.

## Acknowledgements

This work was funded by the Innovative Young Biotechnologist Award (IYBA) 2010, Department of Biotechnology, Ministry of Science and Technology, Government of India.

